# SpaTM: Topic Models for Inferring Spatially Informed Transcriptional Programs

**DOI:** 10.1101/2025.01.24.634726

**Authors:** Adrien Osakwe, Wenqi Dong, Qihuang Zhang, Robert Sladek, Yue Li

**Affiliations:** Quantitative Life Sciences Program, McGill University, QC, Canada; School of Computer Science, McGill University, QC, Canada; Department of Epidemiology, Biostatistics and Occupational Health, McGill University, QC, Canada; Department of Human Genetics, McGill University, QC, Canada

**Keywords:** Spatial Transcriptomics, Topic Modelling, Bayesian Inference, Spatial Segmentation, Neighbour Prediction

## Abstract

**Background:** Spatial transcriptomics has revolutionized our ability to characterize tissues and diseases by contextualizing gene expression with spatial organization. Current spatial transcriptomics pipelines require researchers to use a variety of models and tools to explore spatial domains. However, few methods provide researchers with a way to jointly analyze spatial data from both annotation-free and annotation-guided perspectives using consistent inductive biases and levels of interpretability. A single framework with consistent inductive biases ensures coherence and transferability across tasks, reducing the risks of conflicting assumptions.

**Results:** We propose the Spatial Topic Model (SpaTM), a topic-modelling framework capable of annotation-guided and annotation-free analysis of spatial transcriptomics data. SpaTM can be used to learn gene programs that represent histology-based annotations while providing researchers with the ability to infer spatial domains with an annotation-free approach if manual annotations are limited or noisy. Our benchmarking experiments reveal SpaTM’s competitiveness at spatial label prediction and clustering when compared to state-of-the-art methods. We also demonstrate SpaTM’s interpretability with its use of topic mixtures to represent cell states and transcriptional programs in dorsolateral prefrontal cortex and ductal carcinoma samples and how its intuitive framework facilitates the integration of annotation-guided and annotation-free analyses of spatial data with downstream analyses. Finally, we demonstrate how SpaTM can be used to extend the analysis of large-scale snRNA-seq atlases with the inference of cell proximity and spatial annotations in human brains with Major Depressive Disorder.

**Conclusions:** SpaTM provides researchers with a unified analysis framework for spatial transcriptomics data. By enabling competitive performance in a variety of tasks, SpaTM helps researchers undertake biologically-driven analyses through the identification of interpretable and biologically informed gene programs.

## 1 Background

Advances in Spatial Transcriptomics (ST) have made it possible to reliably profile spatially oriented gene expression throughout the transcriptome. The resulting datasets have enabled novel discoveries by helping researchers identify transcriptionally distinct spatial regions in different tissues and diseases [1, 2]. Many ST analyses have been done in brain and cancer datasets, where researchers have made extensive use of spatial information to compare changes in spatial regions, cell type composition and cell-cell interactions [3–6].

One of the main analyses done for ST is region segmentation [1], which is an initial step in addressing the challenge of exploring domain relationships and identity changes at bordering regions. Segmentation can be performed by training a machine learning model with manual annotations based on on histology- and cytoarchitecture (e.g., Cell Location recovEry or CeLEry) or using spatial clustering approaches (e.g., BayesSpace and SpaGCN) [3, 7, 8]. Although these methods have demonstrated state-of-the-art performance in their respective tasks, there is still a need for methods which can complement these tasks with intuitive, easy-to-interpret results. For example, BayesSpace and SpaGCN identify spatially-resolved markers by first generating spatial clusters and running a differential expression analysis on the clusters. As previously demonstrated by Neufeld et al., such approaches are prone to generating false positives during marker gene detection [9]. Thus, it is preferable to identify methods that jointly learn spatial domains and their markers. Similar observations can be made with supervised approaches that do not guarantee to provide users with an interpretable way to identify the gene programs that drive histology annotations. Finally, most methods are designed to perform only one of these tasks, requiring researchers to integrate results from different methods to perform more comprehensive analyses. This raises new complications, as methods vary in their inductive biases and may not provide results with similar levels of interpretability.

In addition, the limited availability of spatial datasets for specific tissues and diseases requires researchers to find ways to impute spatial information into single cell atlases to perform large-scale spatial analyses [10]. These imputation tasks facilitate our ability to link the findings made in single-cell studies with spatial studies, leading to more nuanced hypothesis generation for experimental efforts. For example, the brains of patients suffering from Major Depressive Disorder (MDD) have been shown to experience distinct morphological changes [11]. Although large-scale single cell analyses of MDD have revealed transcriptional changes in MDD onset, it is still necessary to integrate these findings with spatial information to characterize pathogenetic mechanisms [12]. Thus, finding methods that can reliably model spatial transcriptomes data and project them onto single-cell datasets is crucial to integrate the findings from both modalities.

Currently, there is a lack of an interpretable computational framework that can easily be implemented for different types of spatial analyses and impute spatial information while ensuring seamless and intuitive integration of downstream findings. Such a framework will greatly aid the analysis of single cell disease atlases. To this end, we present the Spatial Topic Model (SpaTM), an interpretable topic modelling framework capable of spatial deconvolution, spatial label prediction, spot proximity prediction, and de novo inference of spatial gene signatures (Fig. 1). By using the interpretable and flexible framework of the topic model as its foundation, SpaTM is able to easily undertake different forms of spatial analyses with the same inductive bias. This ensures a practical analysis framework while preserving competitive performance in different tasks. Finally, we demonstrate that the SpaTM framework can be used to impute spatial information into a publicly available MDD snRNA-seq dataset to investigate spatially oriented changes in MDD and uncover novel pathogenetic insights.

**Fig. 1.**
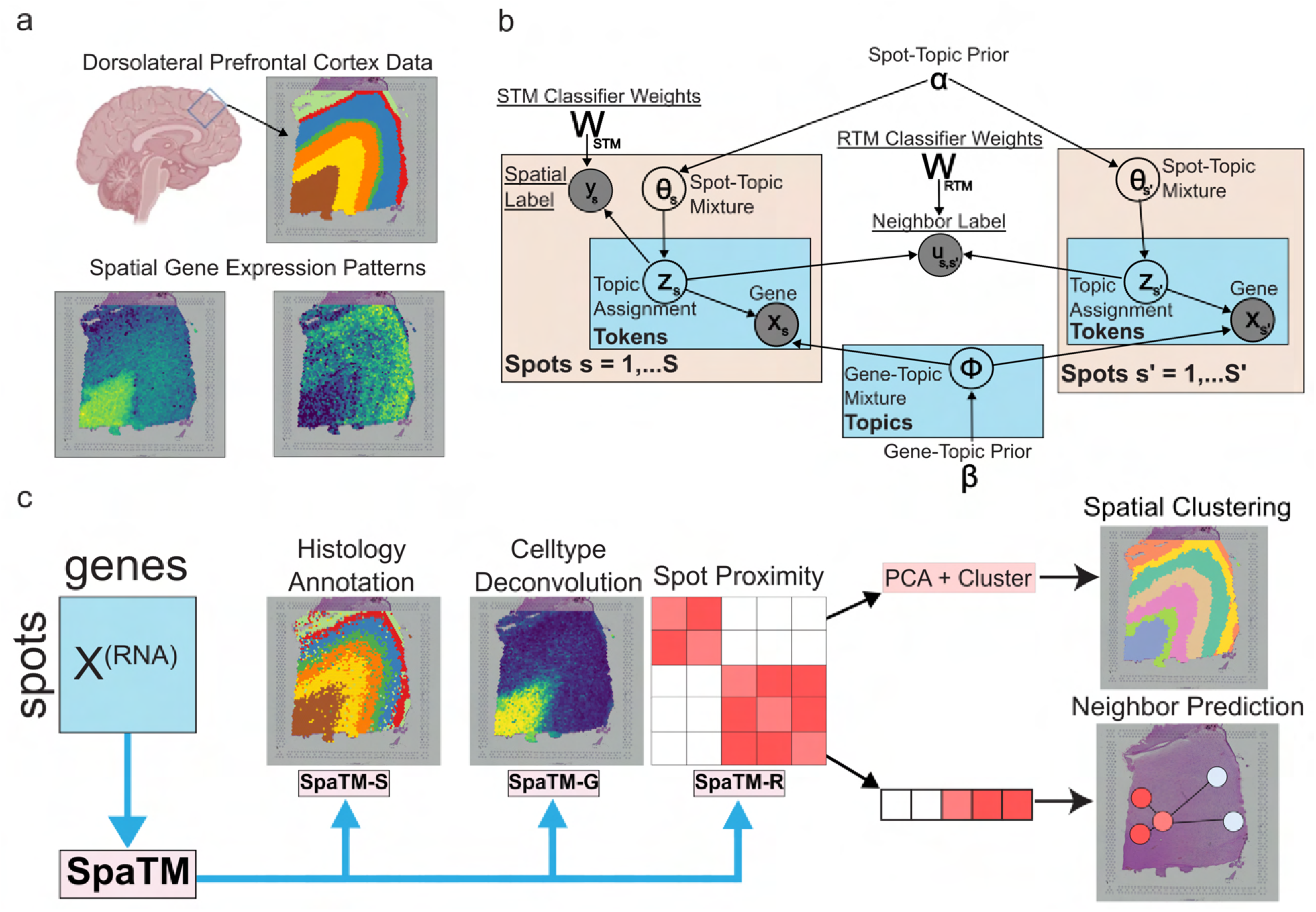
SpaTM for a unified analysis of ST data. **a** Spatial gene expression. Example of a 10X Visium dataset for the dorso-lateral prefrontal cortex containing manual histology-based annotations and spatially-driven gene expression (spatialLIBD). **b** Spatial topic model (SpaTM) modelling framework for ST data depicted as a probabilistic graphical model. The framework extends the GTM-Decon tool to include a supervised topic model (STM) component as well as relational topic model (RTM) component to model spot proximity. **c** Downstream analysis tasks. SpaTM provides a unified framework for ST tasks through its use of the STM (SpaTM-S), GTM (SpaTM-G) and RTM (SpaTM-R) components.

## 2 Results

### 2.1 SpaTM overview

SpaTM is built around the topic model, an interpretable and principled machine learning framework designed to discover latent topics from an unlabelled corpus of documents treated as bags of words (Fig. 1) [13]. The topics are distinct sets of multinomial distributions over the entire vocabulary. In genomics, the topics can be interpreted as modules of co-expressed genes. Due to the natural probabilistic interpretation and the existence of efficient inference algorithms, topic models have a broad and long history of success in many domains that involve sparse and high-dimensional data including scRNA-seq and ST data [5, 14, 15]. In our context, “documents” are spots in ST (or cells in scRNA-seq); “words” represent genes detected by sequencing; “tokens” are individual transcript reads of the same gene and “topics” are biological processes. The value of topic modelling has previously been demonstrated with our Guided Topic Model for deconvolution (GTM-Decon) tasks, which allowed researchers to ensure that the topics learned are anchored to cell types of interest [15]. SpaTM extends GTM-Decon (hereby referred to as SpaTM-G) with two additional components: SpaTM-S for spatial label prediction via supervised topic modelling [16] and SpaTM-R for cell proximity (probability of a pair of cells being neighbours) inference via relational topic modelling (Fig. 1 c) [17]. Thus, this work introduces a novel tool capable of a multitude of ST analysis tasks with high interpretability through a single framework.

### 2.2 Benchmarking SpaTM-S

We evaluated SpaTM-S for the prediction of spatial cortical layers (L1-L6 and white matter) in slices of human dorsolateral prefrontal cortex (DLPFC) using the SpatialLIBD 10x Visium dataset [2]. Benchmarking against state-of-the-art methods involved four scenarios: training on one slice and predicting on another from the same patient (scenario 1) or a different patient (scenario 2), and training on three slices to predict on the same test slices (scenarios 3 and 4 respectively). We compared SpaTM-S’s performance to the results reported by [3] for the same scenarios as well as sequentially training a topic model (i.e. cell type guided with single nuclei reference data from [18] and unguided as the standard topic model) and a *post hoc* classifier (Fig. 2a). SpaTM-S accurately predicts spatial labels from spot-based gene expression profiles and outperforms existing methods for prediction within and between samples (Fig. 2a,b) [3, 19–21]. Updating SpaTM-S’s spatial label predictions based on neighbouring spots via KNN-based majority voting (Smooth) further improved its predictive performance. As our model was trained on spot-resolution data, we assessed its transferability to single-cell resolution by imputing labels on a single-cell DLPFC human sample (Slide-Tags) (Supp. Fig. 1). This revealed that the predicted layers retained spatial coherence in single-cell data. An ablation study showed that SpaTM-S outperforms sequentially training a topic model and classifier (Supp. Fig. 2 A–C). Together, these results demonstrate the superior performance of SpaTM-S in imputing spatial labels over existing methods.

**Fig. 2.**
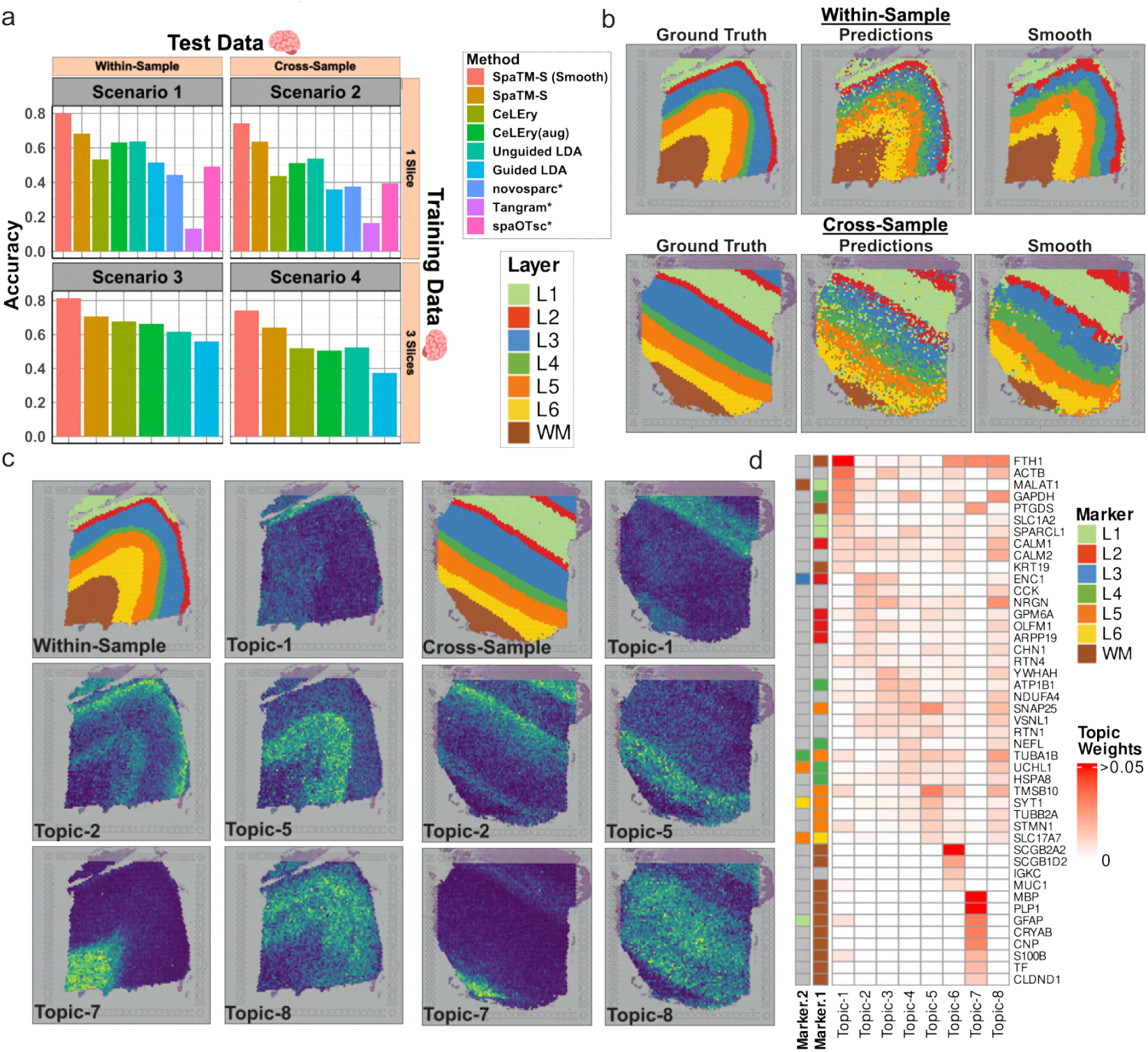
SpaTM-S enables accurate and interpretable histology annotation predictions. **a** Within-sample and cross-sample DLPFC layer predictions. Models were trained on one slice to predict on another from the same patient (Scenario 1) and different patient (Scenario 2), and were separately trained on three slices to predict on the same two test slices (Scenarios 3 and 4 respectively). Asterisks (*) denote methods which can only be trained on a single tissue slice. Results for SpaTM-S’s label smoothing (Smooth) and CeLEry’s data augmentation procedures (aug) were also included. **b** Visualization of SpaTM-S predictions from scenarios 3 and 4. **c** Visualization of the inferred spot-topic mixtures from scenarios 3 and 4. **d** Gene topics learned by SpaTM-S for scenarios 3 and 4. For each topic, the top 10 genes are displayed. The colour intensity is proportional to the probabilities, with values greater than or equal to 0.05 represented as the maximum intensity. Known layer-specific marker genes are annotated on the left. The Marker.1 (Marker.2) annotation represents the layer with the highest (second highest) differential expression (if any) for the corresponding gene as reported by [2].

### 2.3 SpaTM-S interpretability

A key advantage of the SpaTM-S framework is the ability to identify spatial gene programs using the inferred topic matrices 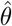 and 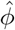. Visualizing the spot-topic mixtures 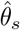 for each spot from unseen histology slices revealed spatial topic patterns that strongly align with the ground-truth layer regions (Fig. 2c). These patterns were present in both the within- and cross-sample scenarios, demonstrating the generalizability of the inferred spatially informed topics. Next, we selected the top genes for each inferred topic in 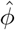 and compared them with the layer-specific markers identified in the SpatialLIBD study. We found considerable overlap, which further supports the accuracy of the spatial gene programs learned by SpaTM-S (Fig. 2d) [2].

### 2.4 Spatially informed gene expression correction

Sparsity and noise in ST data can affect the visibility of spatial gene expression patterns when defining region markers [2]. A solution is to apply pseudo-bulking to spots in different spatial regions to reduce noise in marker expression. However, the loss of spot-level resolution limits our ability to explore how marker expression changes across region borders. To this end, we propose to use the spatially-informed topic mixtures learned by SpaTM-S to correct spatial expression profiles (Fig. 3a). As the topic model factorizes the data into spot-topic and gene-topic mixtures, we can use both metrics to reconstruct the gene expression of a given sample (Fig. 3 a). We trained SpaTM-S on three slices and inferred 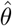 for test slices within and between samples. After reconstructing the spot-by-gene matrix, we scaled the expression profile of each spot by its library size and applied log-normalization. The corrected expression profiles improved the spatial autocorrelation (Moran’s I) of layer markers (identified by pseudo-bulking in [2]) compared to the uncorrected counts (Fig. 3b). The reconstructed profiles also exhibit much clearer spatial patterns (Fig. 3C). To ensure that the reconstruction process was not creating new, unrelated signals, we assessed the correlation between the uncorrected and reconstructed counts with different numbers of genes (Supp. Fig. 2 d-e). We found that both the baseline Latent Dirichlet Allocation (LDA) and SpaTM-S yielded high correlations, suggesting that the de-noised expression is biologically relevant. We also observed similar performance for SpaTM-S and LDA at various gene counts (Supp. Fig. 2 f-g), suggesting the robustness of the topic models.

**Fig. 3.**
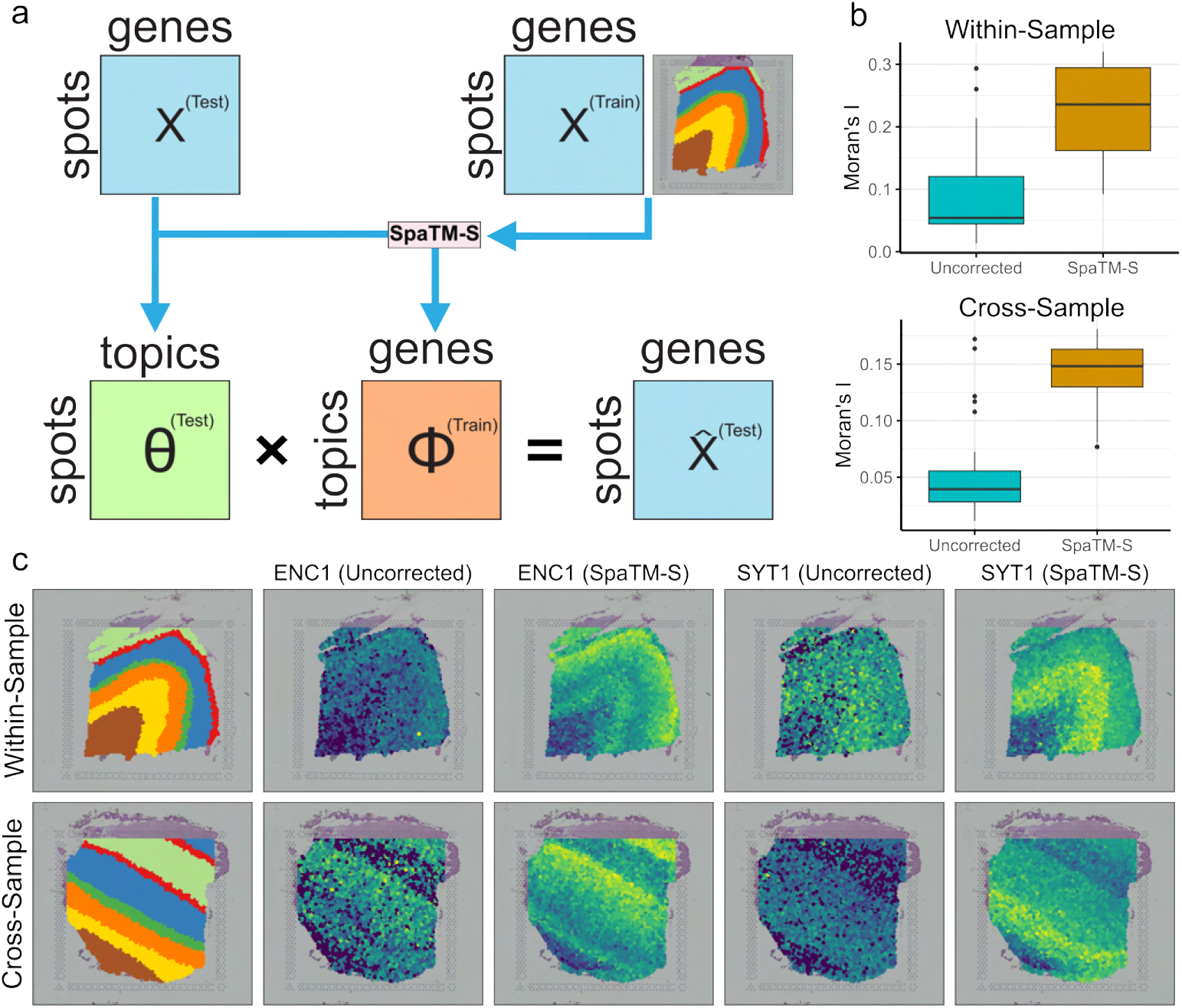
SpaTM-S enables spatially informed gene expression correction. **a** Spot-level gene expression can be corrected with SpaTM-S by taking the matrix product of the inferred spot-topic and gene-topic mixtures. **b** Comparing the spatial autocorrelation (Moran’s I) estimates of previously reported DLPFC markers using log-normalized counts from the uncorrected and SpaTM-S-corrected expression profiles for a within-sample and cross-sample slice. **c** Visualization of spatial expression patterns for DLPFC markers using log-normalized counts from the uncorrected and SpaTM-S-corrected expression profiles.

### 2.5 Benchmarking SpaTM-R

SpaTM-R complements SpaTM-S by constructing a spot or cell proximity matrix to predict neighbours (Fig. 4a). To evaluate neighbour prediction, we trained SpaTM-R in four scenarios: training on one slice and predicting on another from the same patient (scenario 1) or a different patient (scenario 2), and training on three slices to predict on the same test slices (scenarios 3 and 4 respectively). Neighbours were defined as any spot within the radius of a given Euclidean distance. For benchmarking purposes, we used a distance threshold, which provided approximately 10-12 neighbours for each spot. To ensure a balanced training and testing dataset, we then sampled an equal number of spots beyond the threshold distance to use as negative examples. We set our sampling procedure to prioritize the spots that were further from the current spot *s*. We then compared SpaTM-R’s performance to running a logistic regression and 1-layer MLP on the topic mixtures generated from LDA. The evaluation of neighbour prediction for the generated positive and negative examples was performed using AUROC, which revealed the superior performance of SpaTM-R (Fig. 4b). To determine how SpaTM-R handles predicting neighbours within and across spatial regions, we compared the neighbour prediction for within-region and cross-region pairs of spots over different Euclidean distances (Fig. 4c). This revealed that SpaTM-R was more inclined to predict same-region pairs of spots as neighbours than cross-region pairs as their distance increased, suggesting that the neighbour predictions by SpaTM-R capture functional similarity rather than mere spatial proximity.

**Fig. 4.**
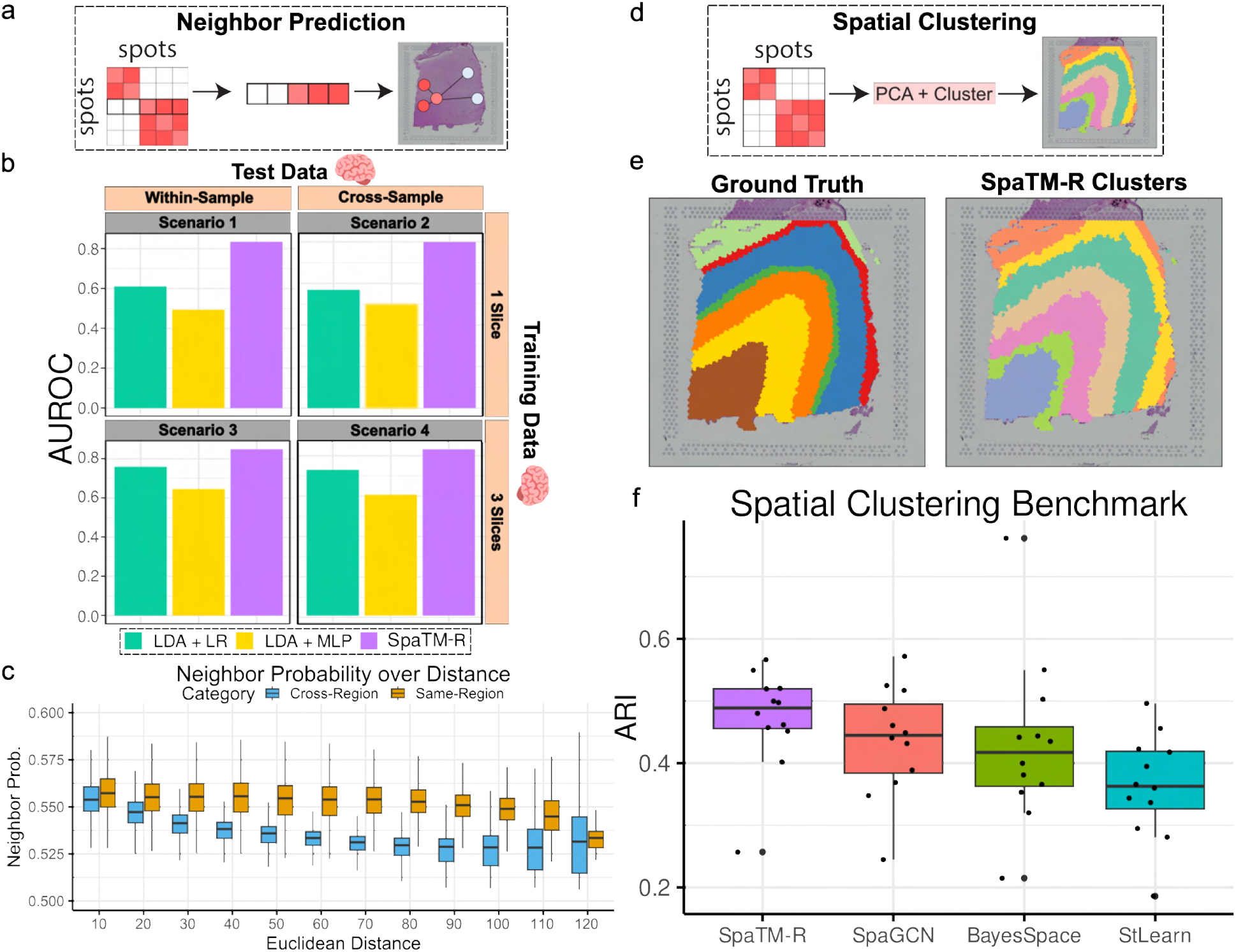
Training SpaTM-R for neighbour prediction yields reliable performance and high-quality spatial clustering. **a** We train SpaTM-R to predict neighbours such that we can infer a spot proximity matrix and neighbours in unseen samples. **b** Benchmarking results for neighbour prediction for SpaTM-R and independently training the LDA and classifier components. We train on one slice and predict on another from the same patient (Scenario 1) and different patient (Scenario 2), and separately train on three slices to predict on the same test slices (Scenarios 3 and 4). **c** Comparison of neighbour predictions for pairs of spots in the same and different regions over different Euclidean distances.**d** SpaTM-R uses the predicted spot proximity to generate spatial clusters. **e** Visualization of smoothened SpaTM-R spatial clustering results for one DLPFC slice. **f** ARI benchmarking for SpaTM-R against state-of-the-art methods on the SpatialLIBD dataset.

Thus, we evaluated the ability of SpaTM-R to generate spatial clusters when manual annotations were unavailable (Fig. 4d). Specifically, we trained SpaTM-R to predict neighbours in each slice from the Spatial-LIBD dataset and used the predicted spot proximity matrices as inputs to Principal Component Analysis (PCA) followed by Leiden clustering. A visual inspection of the generated clusters reveals their close resemblance to the ground-truth layer labels (Fig. 4e). We then compared these results to previously reported benchmarks for state-of-the-art methods on the same 12 DLPFC slices [8]. SpaTM-R demonstrated competitive performance to existing methods while retaining its interpretable framework (Fig. 4f). We also compared SpaTM-R’s clustering performance with a previously published benchmarking study of 14 methods on 34 spatial transcriptome samples from a variety of spatial technologies (spot and single cell resolution) and tissues [22]. Evaluating cluster accuracy with NMI, HOM and COM further highlighted SpaTM-R’s competitiveness vis-à-vis existing tools (Supp. Fig. 3).

### 2.6 Integrative analysis of DLPFC with the SpaTM suite reveals spatial sub-regions

A crucial advantage of SpaTM is its ability to perform multiple tasks and easily integrate their results. The highly interpretable SpaTM suite makes the integration of results from its different tasks more reliable and intuitive than using a pipeline of independent methods. We demonstrate this using one DLPFC slice from the SpatialLIBD database to integrate results found using the SpaTM-S, SpaTM-R and SpaTM-G components. This revealed that white matter spots (as determined by SpaTM-S) were represented by two clusters found by SpaTM-R (clusters 2 and 4) (Fig. 5a). We found that cluster 4 represented white matter spots bordering layer 6 whereas cluster 2 represented the core white matter region. Comparing the SpaTM-S-derived topic mixtures between white matter spots in both clusters shows a clear decrease in the white matter-associated topic weight (Fig. 5b). Finally, running a cell-type deconvolution analysis by training SpaTM-G on healthy DLPFC samples from a reference snRNA-seq dataset revealed that the core white matter cluster had a higher proportion of oligodendrocytes, while the border cluster revealed the appearance of excitatory neurons (Fig. 5c) [12]. As these observations were made on a single slice, we repeated the analysis on three additional DLPFC slices from the same patient. This analysis revealed similar findings with the white matter regions across all four samples exhibiting a drop in oligodendrocyte proportion and an increase in excitatory neurons (Supp. Fig. 4).

**Fig. 5.**
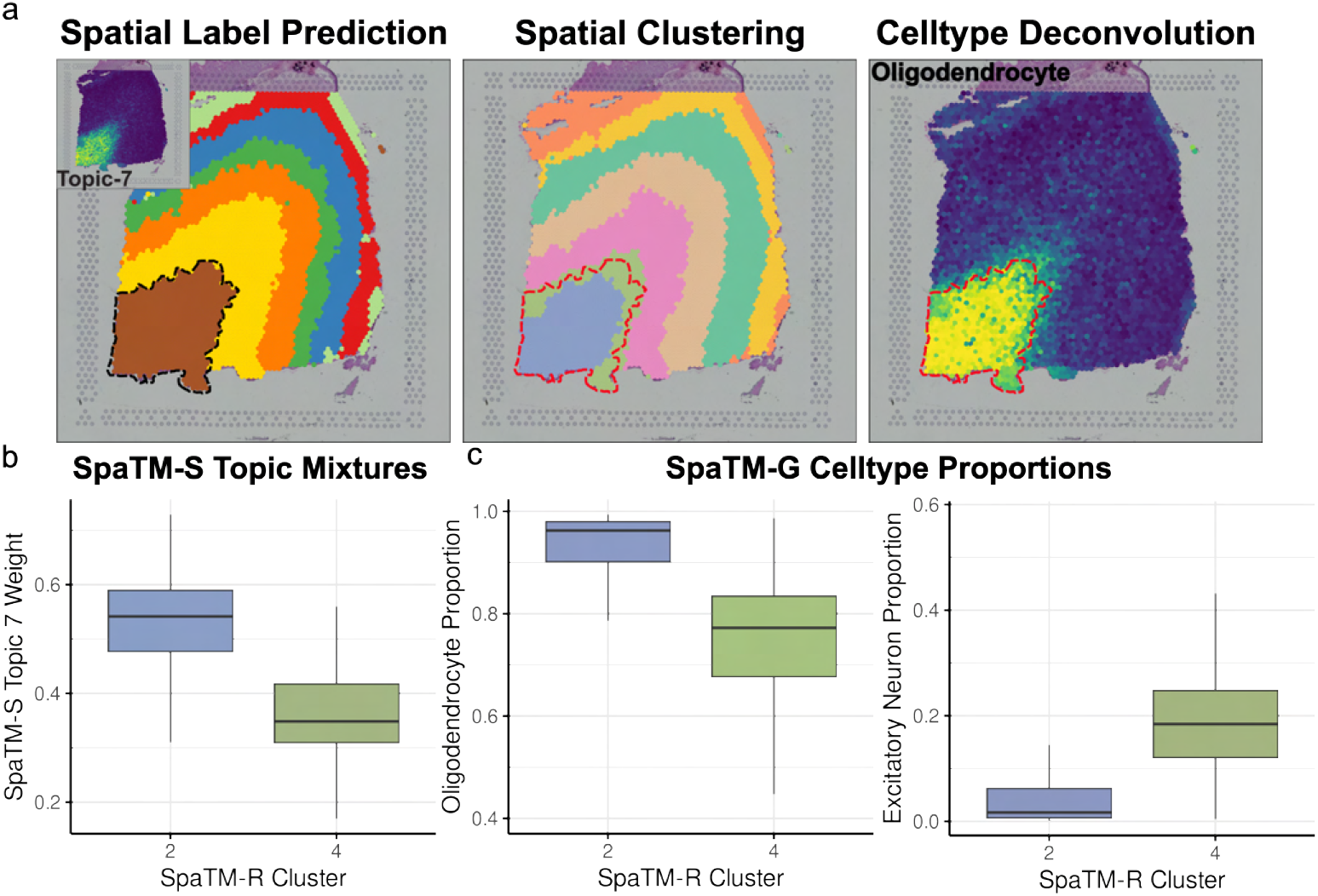
SpaTM enables efficient and intuitive integration of spatial analysis results. **a** Results from the SpaTM suite for one DLPFC slice. SpaTM-S (left) provides the spatial label predictions, SpaTM-R (centre) provides spatial clusters and SpaTM-G (right) provides cell type proportions (visualization shows oligodendrocytes). The dotted line delineates the predicted white matter spots. **b** Comparison of SpaTM-S-derived topic proportions across white matter sub-regions defined by SpaTM-R. **c** Cell-type Deconvolution results (SpaTM-G) comparing cell type proportions in SpaTM-R white matter sub-regions using SpaTM-G.

Together, our proposed SpaTM suite has greatly facilitated integrating results from all three tasks and has identified that the predicted white matter region (as predicted by SpaTM-S) is composed of two domains (as predicted by SpaTM-R) that harbour distinct cell-type proportions (as predicted by SpaTM-G).

### 2.7 Imputing spatial labels and cell proximity for MDD snRNA-seq

In addition to providing a flexible analysis toolkit for ST datasets, SpaTM can be used to impute spatial information into single-cell atlases, where for many diseases, large-scale ST datasets are not available. We acquired a public snRNA-seq dataset from the DLPFC of post-mortem human brains diagnosed with MDD or otherwise healthy [12]. The dataset comprised 160k nuclei from 71 patients, where male and female samples were collected in separate batches. Here, we applied both the SpaTM-S and SpaTM-R models, trained with the above SpatialLIBD data, to impute spatial labels and cell proximity information within the atlas (Fig. 6a). Analyzing the imputed cell proximity matrix revealed strong clustering by both cell types and imputed layers, demonstrating the reliability and coherence between the SpaTM-S and SpaTM-R imputations (Fig. 6b). Jointly ordering the cell proximity matrix by layers and cell type reveals that imputed layers are largely composed of biologically relevant cell types and retain inter-layer cell-type-specific patterns in both control and MDD samples. As male and female samples were collected in different batches, we ran a sex-specific analysis to compare differential layer proportions at a cell-type-specific level (Fig. 6c). In women, we found that MDD cases had a higher proportion of oligodendrocytes (Oli) and oligodendrocyte progenitor cells (OPC) in layers 1 (L1) and 3 (L3), respectively. We also observed a shift of inhibitory neurons (InN) from layer 3 (L3) to layer 2 (L2). Finally, both males and females with MDD exhibited a higher proportion of microglia (Mic) in layer 2 than control samples. A Mann-Whitney *U* test on all combinations of layers and cell types revealed the above results to be significant. In general, MDD samples exhibited a higher level of variability in layer proportions compared to healthy controls. These findings demonstrate the added value of SpaTM through its ability to enable spatially informed analyses of existing disease atlases.

**Fig. 6.**
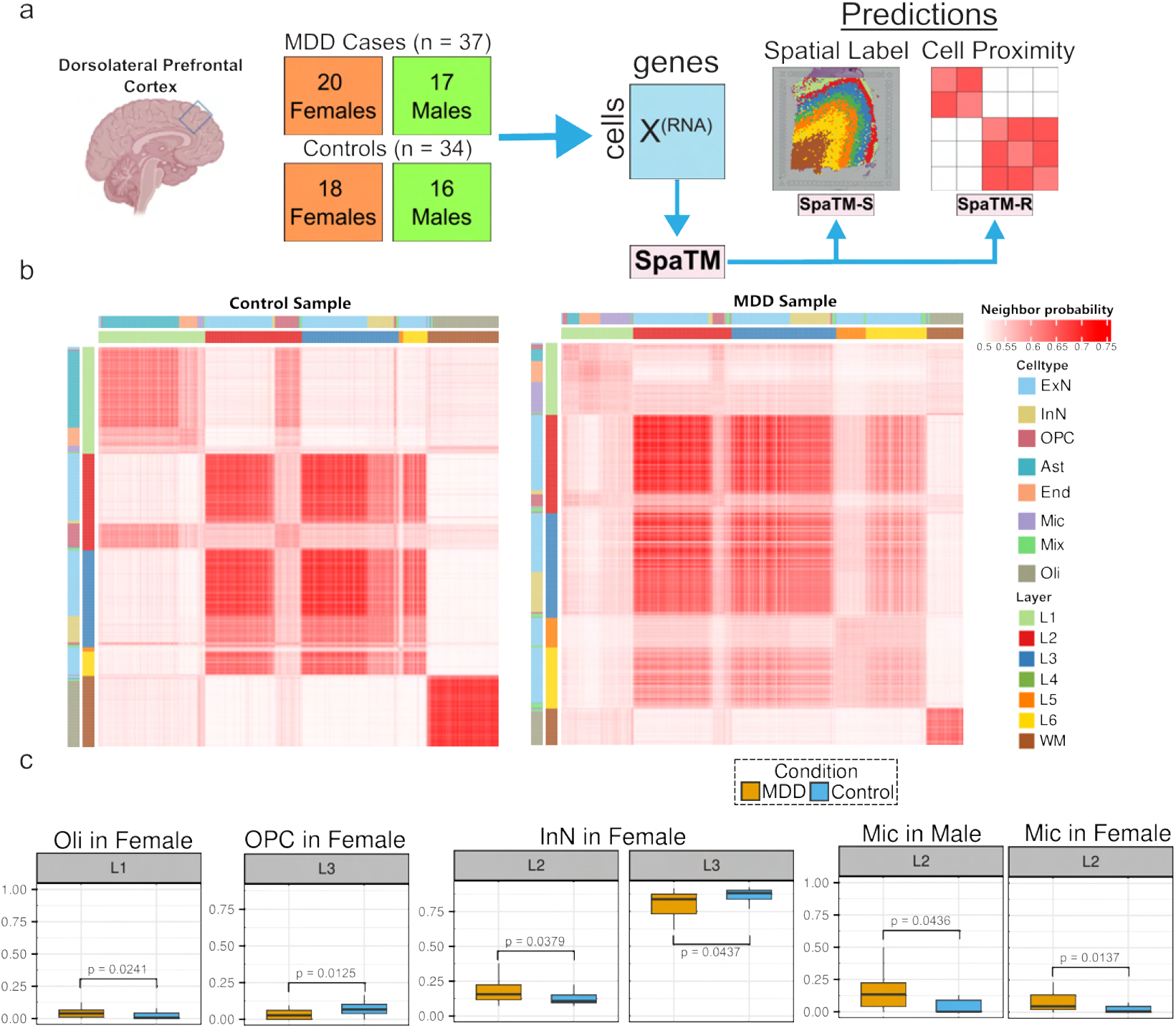
Sex-specific spatial analysis of snRNA-seq MDD using SpaTM spatial imputations. **a** Large-scale DLPFC snRNA-seq atlas for MDD. We use SpaTM to impute spatial labels (SpaTM-S) and cell proximity (SpaTM-R). **b** Heatmap of cell proximity matrix imputed by SpaTM-R for a healthy (left) and MDD (right) female sample. The cells in the heatmap are ordered by cell type nested within SpaTM-S layer predictions. **c** Assessing sex-specific differences in cell type proportions between healthy and MDD samples in different cortical layers using SpaTM-S predictions.

### 2.8 Exploring breast cancer tumour micro-environment with SpaTM-R

We further explore the use of SpaTM on a Visium dataset of breast cancer (ductal carcinoma) [23]. While traditional methods require a downstream differential gene expression analysis to identify cluster-specific markers, SpaTM-R automatically learns cluster-associated markers through its inferred topics. We trained SpaTM-R on a ductal carcinoma ST sample and ran Leiden clustering on the predicted adjacency matrix. The model was trained using 20 topics to match the number of manually annotated sub-regions provided with the dataset [24]. The clusters generated by SpaTM-R show considerable resemblance to the fine pathologist annotations, identifying distant tumour/TME regions with similar expression patterns (Fig. 7a). In particular, we identified a series of topics that represented tumour and micro-environment compartments with distinct gene expression patterns (Fig. 7 b-c). The tumour micro-environment topics incorporated different sets of genes related to anti-tumour and anti-angiogenesis processes. The tumour-specific topics reveal subtypes of invasive and non-invasive ductal carcinoma regions. We further characterized the clusters by running a GSEA analysis using FGSEA on the cancer hallmark pathways on the gene-topic mixtures [25, 26]. Topic 20, a non-invasive region, was enriched for multiple metabolic and cell cycle-associated pathways such as glycolysis, oxidative phosphorylation, the P53 and MTORC1 pathways and androgen/estrogen responses. We also see signs of immune interaction from the enrichment of interferon gamma response and IL2/STAT5 signaling. Topic 11, which represents a specific invasive ductal carcinoma region, was enriched for epithelial-mesenchymal transition (EMT), which supports the tumour region’s progression from non-invasive ductal carcinoma. Both Topics 17 and 19 were enriched for allograft rejection, supporting the TME’s immune response to the proliferation of the tumour region. Both topics are also enriched for EMT, a possible consequence of the interaction with the tumour represented by Topic 20. This case study demonstrates SpaTM’s ability to provide insightful findings on tissues and highly interpretable gene programs that characterize the inferred spatial regions.

**Fig. 7.**
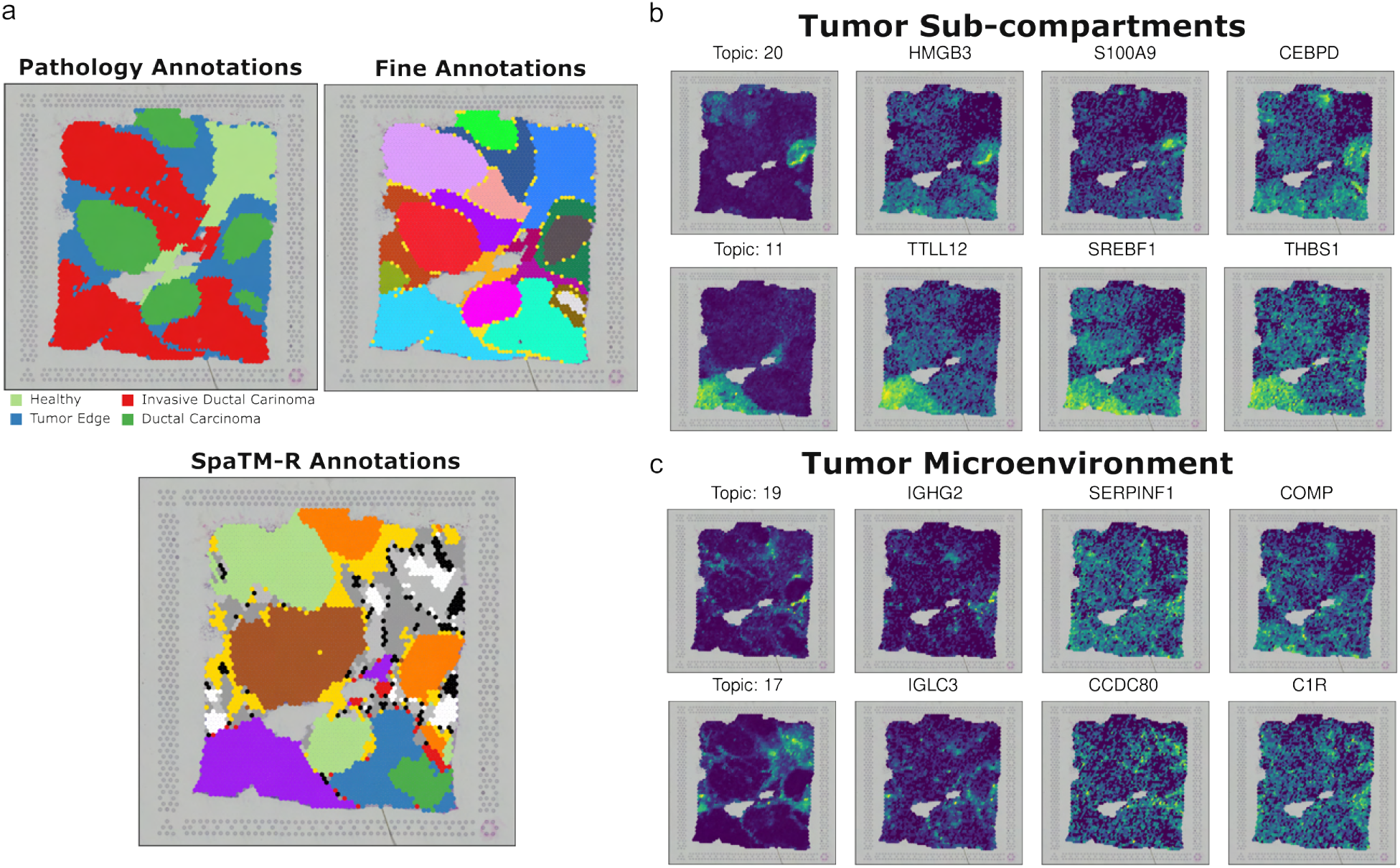
SpaTM-R can characterize tumour heterogeneity in an independent breast cancer dataset. **a** An independent breast cancer Visium sample with broad (left) and fine (right) annotations was analyzed using SpaTM-R. The trained model was used to infer spot proximity and derive unsupervised clusters (bottom) **b** Topics inferred by SpaTM-R during training which identify two distinct tumour regions. **c** Distinct TME regions defined by topics inferred by SpaTM-R.

## 3 Discussion

In this study, we presented a multi-purpose Bayesian topic modelling framework called *SpaTM* to undertake a variety of tasks for ST analyses. Combining guided, supervised, and relational topic model components, SpaTM enables researchers to identify gene programs through interpretable, task-informed topics. Through our analyses and benchmarking studies, we have demonstrated that SpaTM provides a competitive solution for a suite of common ST analysis tasks.

Our benchmarking of supervised histology annotation prediction models demonstrated the superior performance of SpaTM-S (Fig. 2). Most available methods for this task were designed for one-to-one annotation mapping (novosparc,Tangram, and spaOTsc), and are outperformed by MLP-based classifiers such as CeL-Ery which can leverage multiple training samples for training their classifier [3, 19, 20]. By extending the classifier to jointly use a topic-based inference mechanism, we demonstrate how SpaTM-S can take this observation a step further and yield higher prediction performance (Fig. 2a). This observation is complemented by the added interpretability of the topics inferred by SpaTM-S, which provides supervised topics that are representative of the histology annotations (Fig. 2c-d). This combination of features allows SpaTM-S not only to predict histology annotations in unseen samples with greater accuracy, but also to leverage the interpretability of SpaTM-S to better understand gene programs that define the spatial domains defined by histology or pathology experts. Changes in these gene programs can in turn be used to elucidate the cause behind variations in predicted spatial regions.

In addition, SpaTM can correct gene expression to improve detection of spatially variable genes. As seen in previous work, comparing gene expression in different spatial regions often requires pseudo-bulking to mitigate the effect of noise and dropout [2]. However, the gene expression reconstruction capability of SpaTM, acquired through its underlying topic model framework, gives researchers the ability to de-noise spatial data and thus use spot-level resolution to explore spatial patterns and markers in greater detail (Fig. 3).

In the absence of histology annotations, SpaTM-R can instead model spatial proximity. Although many spatial clustering methods are available, most depend on downstream differential gene expression analyses to identify cluster-specific gene markers. As demonstrated by Neufeld et al., this commonly used approach leads to ‘double-dipping’ which has been shown to produce false positives in marker identification [9]. Our analysis demonstrates how the interpretability of SpaTM resolves this issue by directly providing cluster-associated gene programs during training. Thus, SpaTM’s ability to jointly infer gene programs while learning spatial domains provides a conceptually sound solution to avoid double-dipping.

Using the three modelling components provided by SpaTM, we are able to identify and characterize white matter sub-regions by integrating the inferred predictions and topic mixtures (Fig. 5, Supp. Fig. 4). The ability to rapidly undertake and integrate multiple tasks presents SpaTM as a practical solution to researchers aiming to perform multi-faceted analyses in ST data.

Our analysis of the MDD snRNA-seq atlas reveals how spatial imputation can extend single-cell analyses (Fig. 6). The existing literature suggests that the observed shifts in the proportions of the cell type-specific layer may be related to neuronal damage, with the depletion of OPCs suggested to contribute to depressive-like behaviour [27]. These findings demonstrate how spatial imputation can provide a clearer explanation for MDD by integrating spatially driven changes with gene expression.

Furthermore, our case study of an invasive ductal carcinoma sample identified distinct tumour and TME sub-compartments with corresponding gene programs (Fig. 7). These inferred gene programs may play a pivotal role in helping researchers define new, expression-based spatial regions in tissues where histology and pathology expertise are limited.

## 4 Conclusions

Advancements in ST analysis workflows depend on the availability of tools that can streamline and integrate multiple complementary tasks with reliable performance and interpretability. SpaTM addresses this necessity by providing researchers with a flexible and interpretable analysis framework for ST. It enables cell-type deconvolution, spatial label prediction, expression correction, cell proximity inference and spatial clustering with a consistent inductive bias. In addition to its quantitative performance, SpaTM surpasses previous methods by enabling rapid inference and interpretation of spatially driven gene programs.

In future work, SpaTM can be extended to infer cell-cell communication from the predicted cell-cell proximity matrix by using bivariate spatial correlation metrics such as Lee’s L from the Voyager package [28]. These approaches could leverage the inferred topics to identify distant and proximal spatial regions within which distinct gene programs are interacting. In addition, methods such as CellAgentChat [4], could provide additional evidence for ligand-receptor pairs driving the spatial domains identified by SpaTM-R.

## 5 Methods

### 5.1 SpaTM-G —— Guided Topic Model

We assume that each spot *s ∈* {1, …, *S*} is a mixture of *K* topics. This mixture of topics *θ*_*s*_ is in turn sampled from a *K*-dimensional Dirichlet distribution *θ*_*s*_ ∼ *Dir*(*α*_**s**_) parameterized by *α*_*s*_. Our guided topic model framework (SpaTM-G) anchors *α*_*s*_ to cell types, enabling cell type deconvolution on unseen samples [15]. For each spot *s*, with a library size *N*_*s*_, the assignment of topics for each read, *z*_*i,s*_, for read *i* ∈ {1, …, *N*_*s*_} is randomly sampled from one of the *K* topics based on the corresponding topic mixture by *z*_*i,s*_ ∼ *Categorical*(*θ*_*s*_). With the topic assignment *z*_*i,s*_ = *k*, we then map the *i*^*th*^ read to a gene *x* based on the distribution of genes for the sampled topic *x*_*i,s*_ ∼ *Categorical*(*ϕ*_*k*_), where *ϕ*_*k*_ is sampled from a Dirichlet distribution over all genes with *ϕ*_*k*_ ∼ *Dir*(*β*). Thus, the key objective of the topic model is to learn a spots-by-topics matrix *θ* containing the mixture of topics representing each spot and a genes-by-topics matrix *ϕ* which represents the gene composition of each topic. Using the Dirichlet-Multinomial conjugacy [29], we can obtain simplified posteriors for the topic and gene assignments:

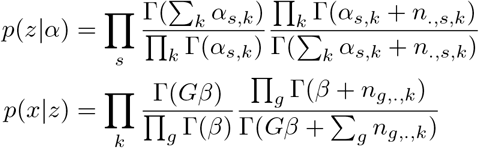

where *G* is the total number of genes in the dataset and the sufficient statistics *n*_.,*s*.*k*_ and *n*_*g*,.,*k*_ represent the number of reads assigned to topic *k* for spot *s* and gene *g*, respectively. This also implies that we can calculate the expected values of *θ*_*s*_ and *ϕ*_*k*_ with the posterior estimates of our topic assignments as they are proportional to the sufficient statistics. The conditional distribution of the topic assignment for read *i* in spot *s*, while fixing the assignment for all other tokens can be represented in closed-form:

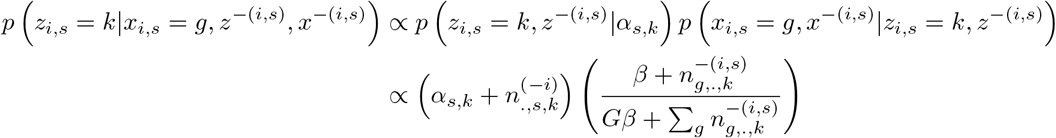

Topic inference can be achieved by collapsed variational Bayes (CVB) [30], where we approximate the posterior topic assignment distribution *p*(*z*_*i,s*_|*x, α*) by 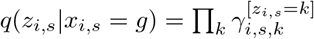 and

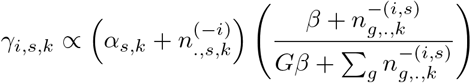

where the sufficient statistics are calculated using the variational parameters as topic assignments:

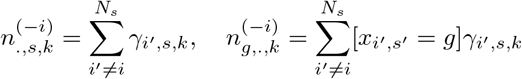

By assuming that all reads for a given gene *g* in a spot *s* have the same topic assignment posterior distribution, we can calculate the variational parameters at the level of genes instead of reads, thereby improving computational efficiency:

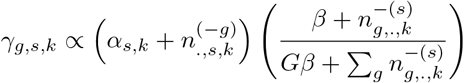

Therefore, we alternate between estimating the variational parameter *γ* and updating the sufficient statistics *n*_.,*s,k*_ and *n*_*g*,.,*k*_. This iterative approach maximizes the evidence lower bound (ELBO) [30]:

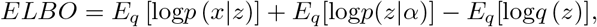

which can assess model convergence. The expected topic mixtures *θ*_*s*_ and *ϕ*_*k*_ can be estimated as:

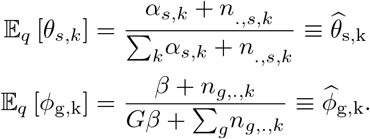

Inference on unseen observations can be achieved by fixing the gene-topic component of the variational parameter estimate to be 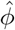 such that

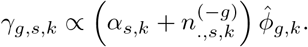

Running the above estimate until convergence can then enable us to infer 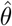 for unseen observations as

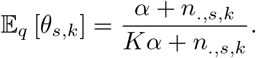

A complete derivation of the base GTM framework and its inherited cell type deconvolution benchmarking can be found in the original GTM-Decon paper [15].

### 5.2 SpaTM-S —— Supervised Topic Model

By using topic mixtures learned by a topic model as input variables for predicting a spatial region *y*, we can reduce high-dimensional and noisy gene expression data to a few interpretable topics that represent spatially-oriented gene programs. Although this can be done by sequentially training LDA and classifier, the topics are not optimized for spatial regions. Thus, we developed a supervised topic model (SpaTM-S), an extension of LDA that jointly trains LDA with a classifier [16] to learn the underlying gene programs (topics) that predict spatial regions. We assume that the spatial label *y*_*s*_ *∈* {1, …, *L*} is generated from a categorical distribution:

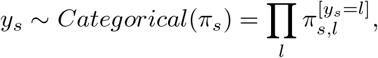

where *π*_*s*_ represents a 1 *× L* vector of probabilities generated by the softmax outputs of a logistic classifier:

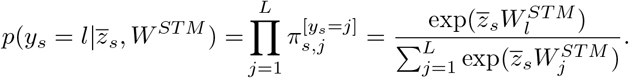

where the weights are represented by a *K* × *L* matrix *W* ^*STM*^ and 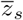 is a 1 *× K* vector that represents the average topic assignment for spot *s*, with the *k*^*th*^ entry of 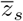 given by

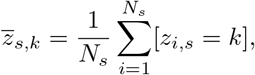

where *N*_*s*_ represents the total number of reads for spot *s*. We can then provide a posterior for the spatial label that fixes all topic assignments but *z*_*i,s*_:

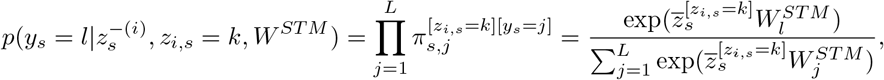

where

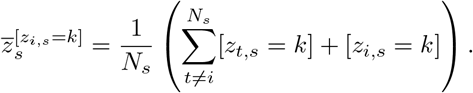

Since we know the ground-truth label for each sample, we can rewrite the above probability such that it represents our confidence in each topic assignment’s ability to predict the label *y*_*s*_

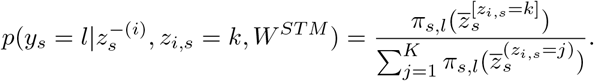

We can thus update the variational parameter estimate to account for spatial label predictions as

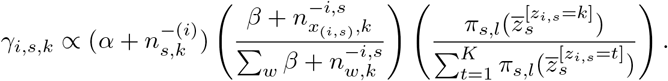

As with SpaTM-G, SpaTM-S can be trained using expectation-maximization by estimating the variational parameter in the E-step and optimizing the classifier weights using updated average topic assignments in the M-step. The model’s convergence is determined using an updated evidence lower-bound

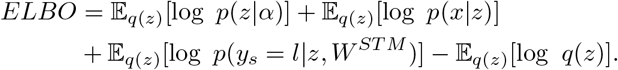

We can also use a multi-layer perceptron (MLP) to predict the labels:

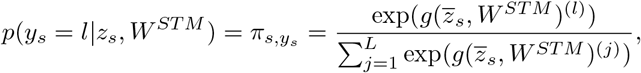

where 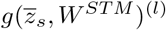 is the output for class *l* for an MLP with parameter weights *W*^*STM*^. After training, we can make predictions on unseen cells by using their topic mixture 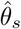 as input to the trained classifier, where 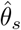 is inferred by the topic model with the gene-topic component fixed to be the estimate 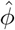 from the training data.

### 5.3 Spatially informed gene expression correction

The gene-topic mixtures 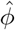 learned by SpaTM-S can be used to reconstruct the expression profiles of unseen samples to reduce sparsity and noise. We can infer the spot-topic mixtures 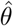 for an unseen slice and compute the matrix product with 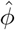 to create a corrected spot-by-gene expression matrix 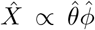. We scale the corrected expression profile of each spot by their library size, making them compatible with existing analysis pipelines. We evaluate the effectiveness of the correction by comparing the spatial autocorrelation in terms of Moran’s *I* of layer markers when using log-normalized counts from the uncorrected and SpaTM-S-corrected profiles:

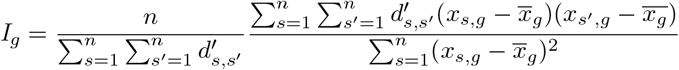

where *n* is the total number of spots for a sample, *x*_*s,g*_ is the expression of gene *g* for spot *s*, 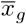 is the average expression for gene *g* and *d*^*′*^_*s,s*_*′* is the inverse of the Euclidean distance between spots *s* and *s*^*′*^ [31].

### 5.4 SpaTM-R —— Relational Topic Model

The relational topic model (RTM) is a variant of STM that was designed to predict links between documents [17]. Here, we adapt RTM to predict spot proximity and neighbours in ST data (SpaTM-R). We assume that for a pair of spots, *s, s*^*′*^, we have a binary neighbour indicator *u*_*s,s*_*′* such that

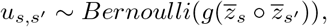

where *g*(*x*) represents a link probability function that defines the probability of spot *s*^*′*^ being a neighbour of spot *s* and 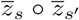 represents the element-wise product of the average topic assignment for each spot. Thus, *g*(*x*) enables SpaTM-R to predict neighbouring spots based on the similarity (or dissimilarity) of their expression profiles. We fix our topic assignments and set our link probability function to be a sigmoid function, where

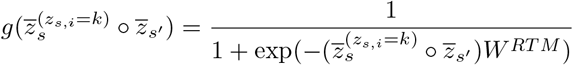

The Bernoulli likelihood for the neighbour predictions of a given spot *s* is then defined as:

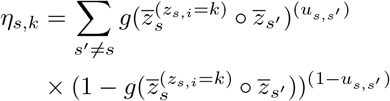

Finally, the topic inference is defined by the joint likelihood of the transcriptome and spatial components of the data:

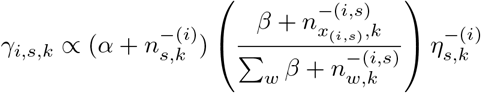

We use Expectation-Maximization to update the variational parameters and optimize a logistic regression model. Convergence is evaluated with the ELBO:

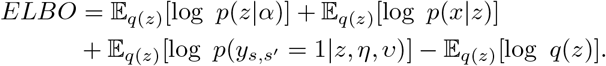

For predictions of pairs of unseen cells *s* and *s*^*′*^, we first infer their topic mixtures 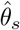 and 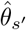, and then use them as input to the trained classifier to predict their proximity.

### 5.5 Training for neighbour prediction from ST data

To train SpaTM-R for spot proximity and neighbour prediction, we define spots as neighbours if they are within a user-defined Euclidean distance from spot *s*. This results in a large class imbalance (i.e., most spots are not a neighbour of any given spot). To address this, we sub-sample the non-neighbour spots, assigning sampling probabilities that are proportional to their distance from the current spot.

### 5.6 Spatial clustering

We evaluated whether SpaTM-R can recapitulate spatial information from gene expression. We first ran PCA on the spot proximity matrix predicted by SpaTM-R followed by the Leiden community detection algorithm to generate clusters. If the number of regions is known, we fine-tune the Leiden resolution parameter to produce the desired number of clusters. For ease of visualization, we smooth the cluster labels by updating the label of any spot based on the label of its neighbouring spots using majority voting.

A more comprehensive spatial clustering benchmark analysis was performed following the protocol proposed by the SDMBench pipeline, where 13 methods were evaluated on 34 unique ST samples [22]. SpaTM-R was run on all 34 samples using the top 10 nearest spots/cells as positive examples for each spot/cell. If samples had more than 3000 genes in their assay, the top 3000 highly variable genes were selected. Otherwise, all genes were used. The number of topics was set to match the number of ground truth annotations present in each sample. We evaluated the NMI, HOM and COM accuracy metrics with respect to the reported results from SDMBench and also compared the average of the three metrics.

### 5.7 Breast cancer ST analysis

Our case study of tumour heterogeneity in breast cancer was performed by running SpaTM-R on a Visium sample from the 10X Genomics database[23]. We set the number of topics to the number of pathologist-derived fine annotations (20) [24] and used the top 5000 highly variable genes. We then assessed which inferred topics correlated with the ground truth annotations and visualized some of the top 10 genes for four of these topics (two tumour-specific topics and two TME topics).

### 5.8 MDD snRNA-seq imputation

For MDD imputation tasks, we trained SpaTM-S on two DLPFC slices using the top 1000 HVGs and eight topics based on its performance on the DLPFC Slide-Tags dataset (Supp. Fig. 1). SpaTM-R was trained on one slice using the top 5000 HVGs. We inferred the topic mixtures for the MDD dataset using both models to then impute spatial labels and cell-cell adjacency for each patient. Spatial labels were used to calculate the layer distribution of each cell type prior to assessing sex-specific changes between MDD and control samples. Statistical testing was performed using the Mann-Whitney U test.

### 5.9 SpaTM software

As a software package, we have implemented SpaTM in R and extended the SingleCellExperiment and SpatialExperiment object classes to better integrate our tool into large-scale analysis pipelines undertaken by life science researchers [32]. The package is hosted on GitHub at https://github.com/li-lab-mcgill/SpaTM.

## 6 Declarations

### 6.1 Ethics approval and consent to participate

Not applicable.

### 6.2 Consent for publication

Not applicable.

### 6.3 Availability of data and materials

All datasets used in this study were acquired from publicly available sources. The 12 DLPFC Visium samples were acquired from the SpatialLIBD study [2]. The DLPFC Slide-tags sample was acquired from the original Slide-tags paper [18]. The MDD snRNA-seq atlas was acquired from Maitra et al. [12]. Finally, the Visium breast cancer sample was acquired from the 10X Genomics database [23].

The SpaTM package and associated documentation can be accessed at https://github.com/li-lab-mcgill/SpaTM. All analysis scripts used in this study can be found at https://github.com/aosakwe/SpaTM_Analysis.

### 6.6 Author contributions

Y.L. conceptualized the study. A.O. implemented the software, conducted the experiments, and wrote the initial draft of the manuscript. Y.L. and R.S. supervised the study. All authors analyzed the results and contributed to writing the final version of the manuscript.

## 6.7 Acknowledgements

We thank Doruk Cakmakci and Anjali Chawla for their helpful feedback on data visualizations and the MDD analysis.

## Appendix A

### Supplementary Figures & Tables

**Supp. Fig. 1.**
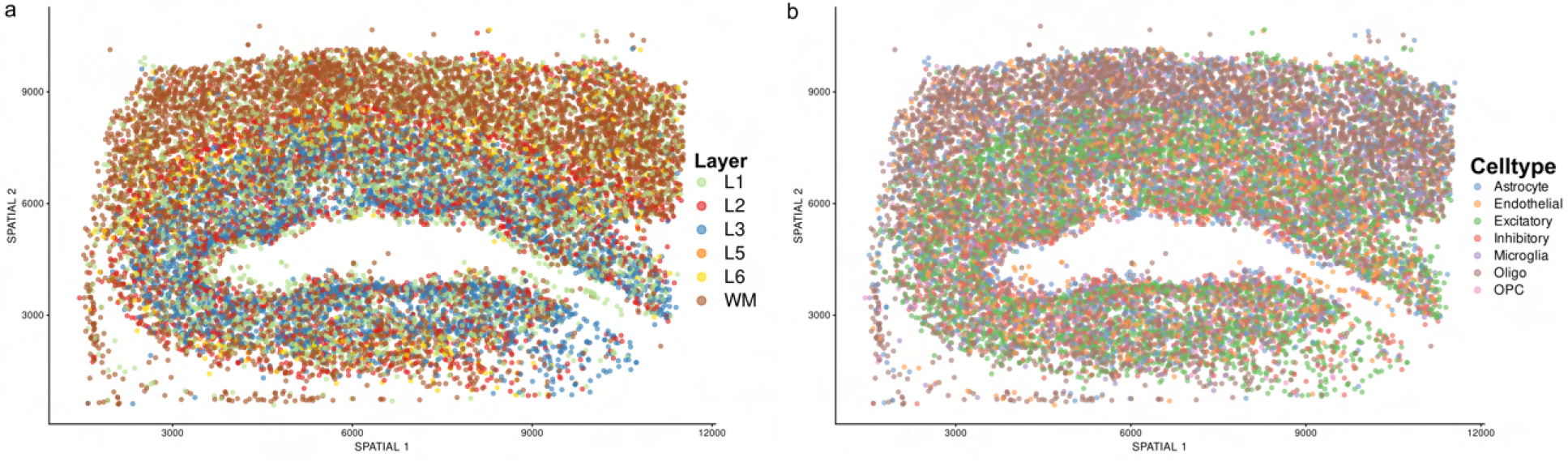
Predicting spatial labels in single cell ST data using spot resolution training data with SpaTM-S. **a** Results from imputing spatial labels with SpaTM-S on a Slide-tags dataset of human DLPFC at single cell resolution after training on spot-resolution data. **b** Spatial distribution of DLPFC cell types.

**Supp. Fig. 2.**
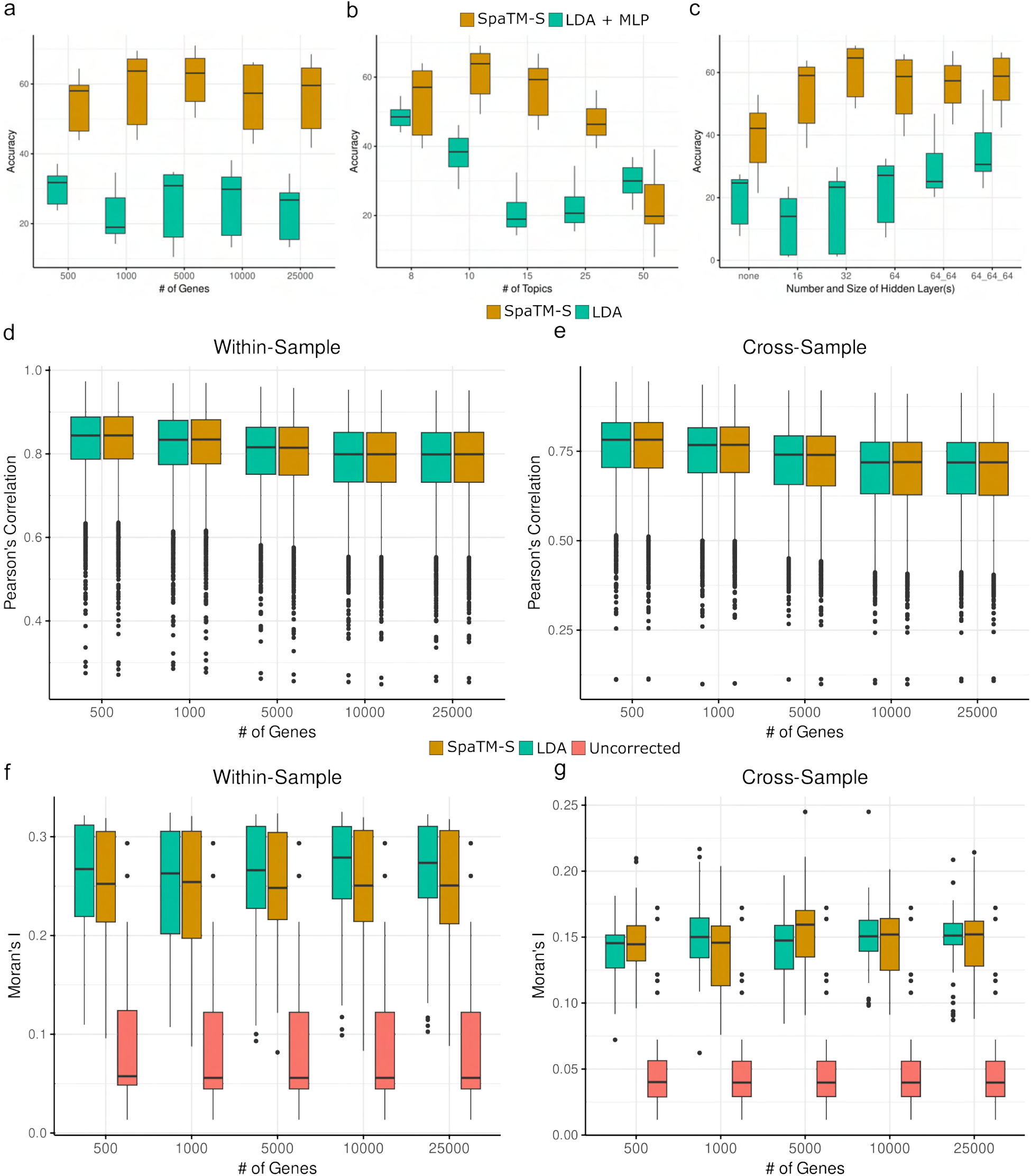
Hyper-parameter Stability of SpaTM-S for label prediction and gene expression correction. **a-c** Ablation study comparing the performance of SpaTM-S at predicting labels in 11 DLPFC samples against using a pipeline approach (LDA + MLP) with different numbers of genes (**a**), topics (**b**) and different classifier architectures (**c**). **d-e** Evaluation of the spot-level correlation of uncorrected and corrected (LDA/SpaTM-S) gene expression after training on a sample from the same (**d**) and a different (**e**) patient. **f-g** Moran’s I of known DLPFC markers using uncorrected and SpaTM-S/LDA-corrected gene expression for a sample from the same (**f**) and a different (**g**) patient.

**Supp. Fig. 3.**
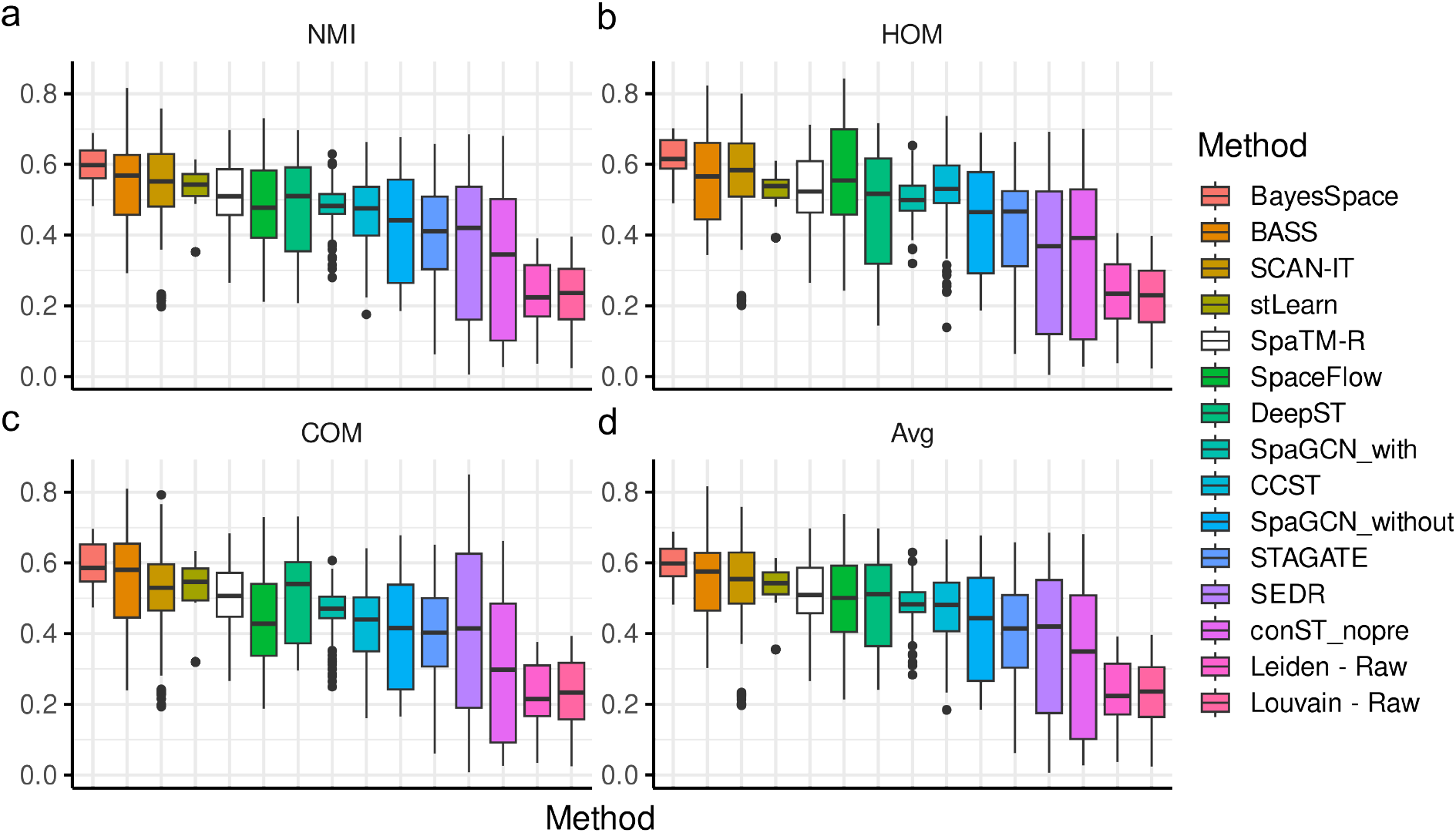
Spatial Clustering benchmarking on multiple methods and ST platforms. Evaluation of spatial clustering methods using NMI **a**, HOM **b** AND COM **c** metrics for 15 different methods on 32 different datasets from different spot-level and single cell spatial technologies. Results for competing methods are as reported in the SDMBench study. **d** Average for the three metrics reported in **a-c**.

**Supp. Fig. 4.**
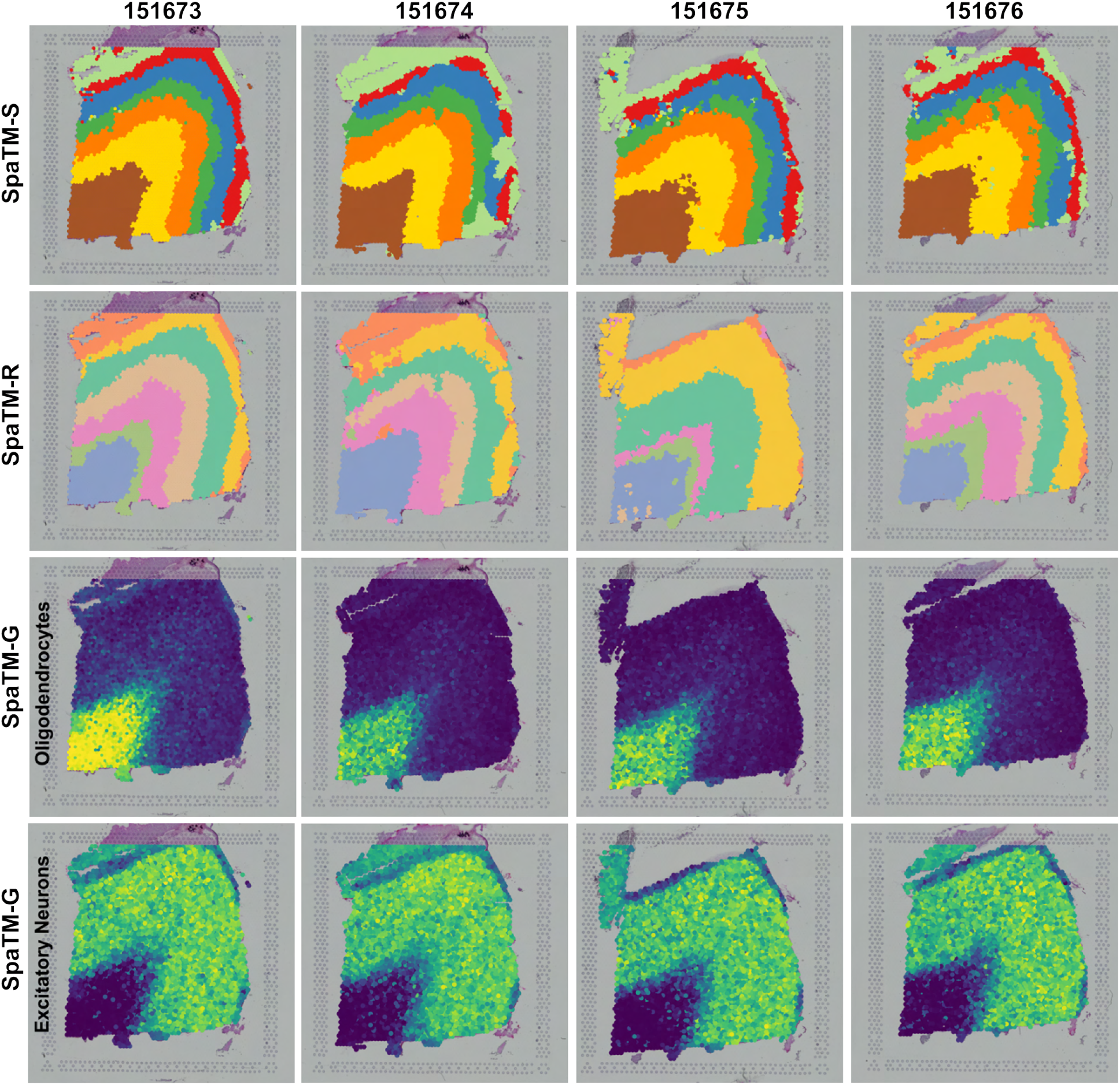
Identifying white matter sub-domains with SpaTM. Results from four DLPFC slices from the same patient integrating spatial label predictions (first row) with annotation-free clustering (second row), oligodendrocyte proportions (third row), and excitatory neuron proportions (fourth row) to identify white matter sub-domains.

